# Prediction of Molecular Mechanisms of Breast Cancer Metastasis

**DOI:** 10.1101/106179

**Authors:** Geoffrey H. Siwo, Gustavo A. Stolovitzky, Solomon Assefa

## Abstract

Metastasis -the spread of cancer to other parts of the body- causes 90% of cancer deaths, underlies major health complications in cancer patients and renders most cancers incurable. Unfortunately, the molecular mechanisms underlying the process are poorly understood and therapeutics to block it remain elusive. Here, we present a computational technique for scanning genome-scale regulatory networks for potential genes associated with metastasis. First, we demonstrate that in the breast cancer cell line MCF7, the commonly dysregulated cancer biomarkers TP53, ERBB2, ESR1 and PGR are closely connected to known metastasis genes with a significant proportion being 2^nd^ degree neighbors of a given biomarker. Next, we identify genes whose 2^nd^ degree neighbors are connected in a similar manner to these biomarkers. Consequently, these are referred to as metastasis associated genes or MAGs. We identify 190 genes that are TP53-MAGs, 22 ERBB2-MAGs, 240 ESR1-MAGs and 84 PGR-MAGs (FDR adjusted *P* <0.001). Analysis of the MAGs reveals statistically significant enrichment with biological functions previously associated with metastasis including the extracellular matrix (ECM) receptor interaction, focal adhesion, cytokine-cytokine receptor interaction and chemokine signaling. The biological significance of MAGs is further supported by their enrichment with experimentally validated binding sites for transcription factors that regulate metastasis, for example BACH1- a master regulator of breast cancer metastasis to bone. The predicted MAGs are also clinically relevant as therapeutic targets for metastasis blocking agents. Specifically, genes that are perturbed by drugs and miRNAs that influence metastasis are enriched with MAGs. Furthermore, some MAGs are associated with patient survival and provide insights into the proclivity for breast cancer subtypes to preferentially spread to specific organs. The results of this study imply that aberrations in primary tumors may constrict metastasis trajectories. This could enable the prediction of organ specific metastases based on aberrations in the primary tumor and lay a foundation for future studies on individualized or personalized models of metastasis. The approach is potentially scalable across other cancers and has clinical implications.

## Introduction

In 1889, Stephen Paget reported the unique dissemination of breast cancer cells in organs of various patients at autopsy leading him to posit that cancer cells are like seeds that spread or metastasize to body parts with compatible soil for their growth^1^. While the ‘seed and soil’ hypothesis has continued to gain support, understanding of the molecular basis of organ specific metastasis remains one of the most vexing questions in cancer research^23–4^. Metastasis is a major clinical challenge in the fight against cancer: it is responsible for over 90% of cancer related deaths, is a leading cause of severe illness in cancer patients, is often associated with multi-drug resistance and leads to rapid death^5^. Metastasis is a hallmark of most solid cancers and many cancer patients eventually develop them. In fact, many patients already have metastases by the time a primary diagnosis is made and on autopsy cancer patients are found to have metastases that were unknown in their life^6^.

The genetic aberrations in the primary tumor may influence organ specific metastases as: i) different cancers show preferential spread to different organs^7,8^, ii) breast cancer subtypes associated with distinct molecular aberrations metastasize with different frequencies to distinct organs^9–14^; iii) some genes involved in cancer initiation that are mutated in the primary tumor such as TP53 also regulate genes that drive metastasis^15–19^, iv) many cancer driver genes are involved in growth while many genes involved in metastasis influence processes such as cell adhesion^20, migration21^ and stem cell differentiation^22, 23^ that are key in developmental processes. Growth and developmental pathways are functionally linked with genes such as TP53 and some growth factors having the ability to influence both growth and differentiation^24,25^.

In breast cancer, the major molecular subtypes of breast cancer namely basal-like type (frequently triple negative breast cancer- estrogen receptor negative- ESR1-, progesterone receptor negative- PGR- and ERBB2 negative- ERBB2-), luminal (A and B) and HER-2 enriched (ERBB2+) subtypes- have distinct prognoses and organ tropisms^13,26–28^. Across all these subtypes, aberrations in TP53 are also highly frequent. TP53 mutations are present in as high as 80% of basal breast cancers^29,30^. Several independent studies indicate that many primary tumors harbor at least some determinants of metastasis ^31–34^. For example, the genomic similarity between primary tumors and their metastasized counterparts from the same patient is much higher than the similarity between primary tumors from different individuals or between metastasized tumors from different individuals ^31,35^. These studies imply that tumor initiating events and other early genomic events may have a molecular link to biological processes that mediate metastasis at later stages. A mechanistic understanding of whether mutations in primary tumors constrict the metastatic landscape is currently lacking yet could be vital for predicting and targeting metastases.

We reasoned that specific cancer biomarkers may be non-randomly associated with metastasis gene networks, thereby increasing the likelihood that aberrations in cancer initiating genes could propagate into the molecular networks of metastasis. These non-random associations may be detected in cancer molecular networks by comparing the network distances between specific cancer biomarkers to those of metastasis driving genes to distances separating the cancer biomarkers to randomly sampled gene sets. As a proof of concept, we used a regulatory network of a breast cancer cell line MCF7, inferred from transcriptional perturbations of the cell line by multiple drugs to examine whether the major cancer biomarkers in breast cancer subtypes are non-randomly associated with metastasis networks.

## Results

### Cancer Biomarkers TP53, ESR1, ERBB2 and PGR are Molecular Neighbors of Metastasis Associated Genes, MAGs

We hypothesized that one way through which early genomic events such as common aberrations in primary tumors may influence metastasis is through close molecular associations between cancer initiating genes and those involved in metastasis. Specifically, genes such as TP53 may be part of biological networks that are close neighbors to those that drive metastasis. Such non-random associations could potentially be inferred from gene interaction networks as an enrichment of known genes associated with metastasis within the network neighborhoods of the biomarkers (Figure 1 A). To test this hypothesis, we obtained a breast cancer interactome constructed using gene expression data from the MCF7 cancer cell line (Methods) and queried the network for the connections between each cancer biomarker and a list of 35 genes linked to metastasis based on various studies 36 (Methods and Supplementary Table 1). We then determined for each biomarker the number of the known metastasis genes located 1 step away from its node in the interactome (direct connections), those located 2 steps away, 3 steps away, 4 steps away and 5 steps away. We then compared the observed number of metastasis genes at each *n* step to that of the same biomarker in 1000 randomly sampled gene sets of the same size as the metastasis gene set (Figure 1 A). This approach controls for the degree connectivity of the biomarker of interest in the input network while preserving the network architecture, thereby providing a measure how close the biomarker is to metastasis genes relative to its closeness to random gene sets.

**Figure 1.**
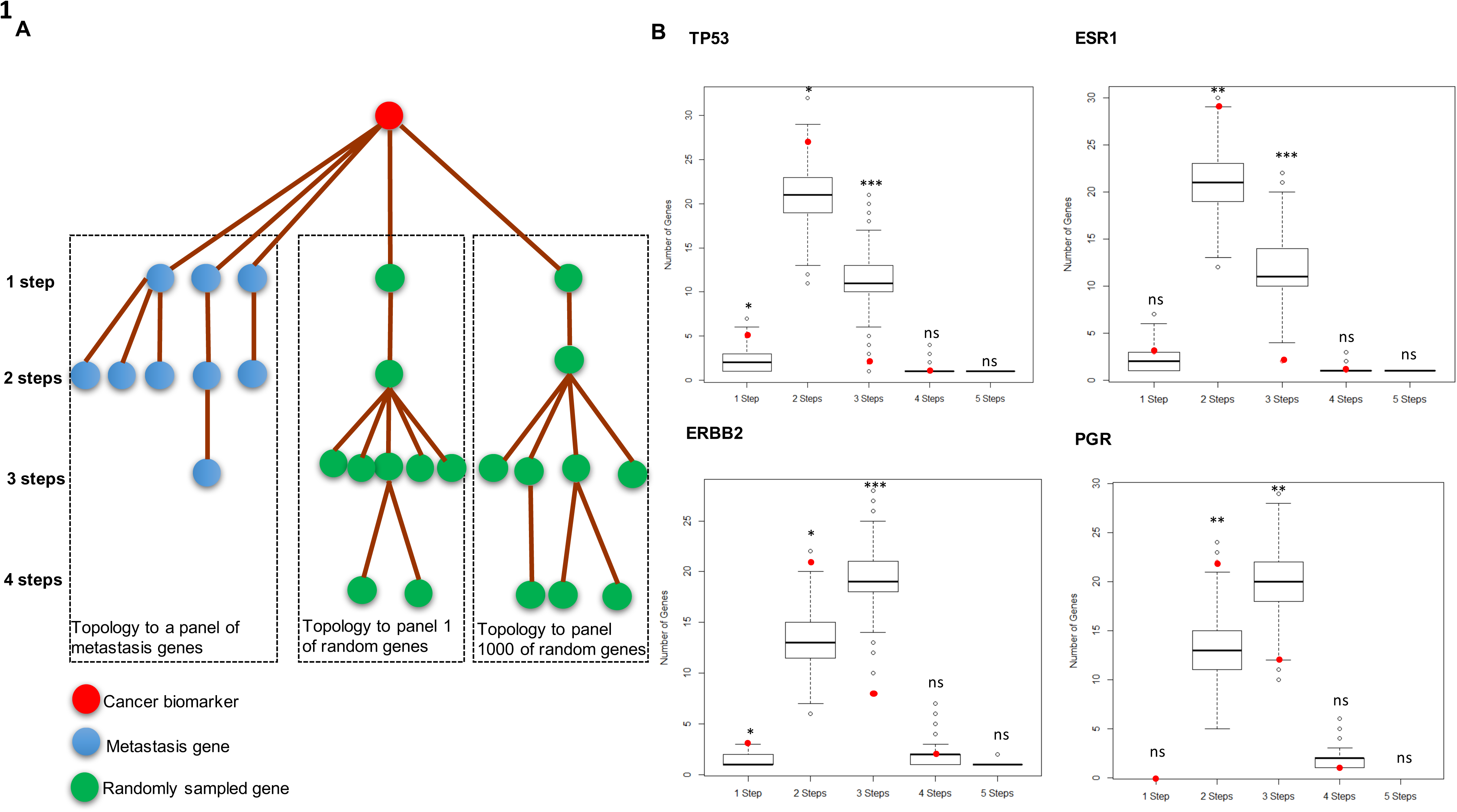
Cancer biomarkers associated with main breast cancer subtypes are non-randomly associated with metastasis genes. (**A**)Traversal of the reverse engineered MCF7 interactome starting from cancer biomarker nodes to previously reported metastasis genes compared to traversal from biomarker nodes to randomly sampled gene sets. (**B**) Number of metastasis genes that are 1,2,3,4 or 5 steps away from a each of the biomarkers TP53, ESR1, ERBB2 or PGR compared to the number of genes in 1000 randomly sampled gene sets at the same steps away from the biomarkers. Highlighted in red is the number of genes in the metastasis gene set that are a given step away from the biomarker specified on each plot. (Significance level: * P ≤ 0.05, ** P ≤ 0.01, *** P ≤ 0.001, ns P > 0.05).

Across all the four biomarkers, we observed significantly more metastasis genes located 2 steps away from the biomarker and significantly less metastasis genes located 3 steps away (*P* < 0.05; Figure 1 B). We examined the 1^st^ and 2^nd^ degree neighbors of TP53 to validate whether they recapitulate some of the previously reported associations between TP53 and metastasis^18^. In the case of TP53, metastasis genes were also enriched in its 1 step or direct neighbors (*P* ~ 0.03) through its direct connections to ERBB2, EGFR, IGFB3, HSPB1 and MUC1 in the interactome (Supplementary Figure 1). These observations are consistent with the role of TP53 in metastasis^16–18^. For example, its inactivation in mouse mammary epithelial cells leads to overexpression of ERBB2^37^ and some of its mutant versions enhance EGFR and ERBB2 signaling^38^, both of which influence metastasis^39^. Furthermore, IGFBP3 is a direct target of wild-type TP53^40^ and expression of IGFBP3 in malignant breast tumors is higher relative to that in adjacent normal tissues^41^. More broadly, TP53 loss can disrupt pathways that inhibit metastasis, defects in TP53 transcriptional activity can lead to gain of functions that promote metastatic processes^18^ and overexpression of the gene is associated with advanced TNM stage and visceral metastases in breast cancer patients^42^. 22 out of 35 of the metastasis associated genes were 2^nd^ degree neighbors of TP53 (*P* ~ 0.01; Supplementary Table 1) including genes involved in cell invasion (matrix metalloproteinases MMP1 and MMP9), chemotaxis (CXCR4) and cell adhesion (CD44, VCAM1, TGFB1 and ITGB6)^36^ (Supplementary Figure 1). ESR1 was 1 step away from 3 metastasis genes (EGFR, MUC1 and USP9X, *P* = 0.17) and 2 steps away from 29 metastasis genes including CXCR4, IL2, IL6, IL8, PTEN and TGFB1 (*P* = 0.002). ERBB2 was directly connected to 3 metastasis genes- EGFR, MUC1 and IL6 (*P* = 0.016; Supplementary Table 1) and 2 steps from 21 metastasis genes including VCAM1, CXCR4, MMP9, MMP2, TGFB1 and CDH1 (*P* = 0.012; Supplementary Table 1). PGR was not directly connected to any of the metastasis genes but 22 of these genes were 2 steps away from its node (*P* = 0.002; Supplementary Table 2 for distances between each of the metastasis genes and the four cancer biomarkers).

The regulatory role of TP53 in metastasis has been an area of active research. Therefore, we assessed the potential regulatory mediators between TP53 and its 2^nd^ degree metastasis genes by examining the intermediary genes linking them in the network (Supplementary Figure 1). These genes encode transcription factors such as SP1, MYC, JUN, RELA, EP300, E2F1 and WT1 that interact with TP53 through protein-DNA interactions or protein-protein interactions (Supplementary Figure 1). Notably, some of these transcription factors have previously been shown to regulate the expression of metastatic processes. For example, SP1 overexpression was shown to inhibit the migration and invasion of breast cancer cells^43^; MYC suppresses the motility, invasiveness and metastatic potential of breast tumors through transcriptional repression of integrins^44^, JUN overexpression enhances liver infiltration of breast cancer cells in nude mice^45^ and RELA inhibits metastasis by transcriptional repression of the breast cancer metastasis suppressor 1 (BRMS1)^46^. We hypothesized that miRNAs that affect expression of these TP53-metastasis intermediary genes could interfere with metastasis. Therefore, we queried miRNA targets (TargetScan^47^) to identify those whose targets are enriched with these genes using Enrichr^48^. mir-29A/29B/29C, mir-507 and mir-7 perturbations had significant enrichment for these genes (*P* < 0.05; Supplementary Material). mir-29s repress cancer metastasis through regulation of the tumor microenvironment^49^ and activate TP53^50^ while mir-7 suppresses brain metastases through its regulatory effect on KLF4 leading to attenuation of cancer stem-like cells invasion and self-renewal^23^. These results validate the ability of the MCF7 interactome to capture known regulatory associations between TP53 and metastasis.

#### Prediction of Metastasis Genes associated with each Biomarker

We reasoned that genes that have the same connectivity as a given cancer biomarker with respect to the metastasis gene set might similarly be associated with metastasis. We defined such genes as those whose 2^nd^ degree neighbors contain a significant proportion of known metastasis genes that are also 2^nd^ degree neighbors of a given biomarker, relative to the proportion of randomly selected genes shared by the biomarker and the gene of interest (Figure 2 A and Methods). To identify such genes with respect to each biomarker, we scanned the whole interactome for additional genes with this topological connectivity to metastasis genes. For TP53, we identified 190 genes as potential metastasis associated genes (MAGs), 240 genes for ESR1, 22 genes for ERBB2 and 84 for PGR (FDR adjusted *P* < 0.001; Figure 2 B and Supplementary Table 3). Functional enrichment analysis of the predicted MAGS (taking into account the size of each of the gene sets) revealed the top enrichments as extracellular matrix (ECM) receptor interaction, focal adhesion, cytokine-cytokine receptor interactions and chemokine signaling (*P* << 0.05; Figure 2 B). We also observed that these functions are enriched only when the high confidence MAGs for each biomarker (*P* < 0.003) are considered, confirming that the observed enrichments are non-random (extended tables for the enrichment analysis are provided in the Supplementary Material). These top functions are all consistent with biological processes that drive metastasis^5,36^ (Figure 2 B), thereby validating that the MAGs are metastasis associated. We compared the overlap between the predicted MAGs for each cancer biomarker to identify genes that are shared across them vs. those that are uniquely associated with a single biomarker. TP53-MAGs were enriched amongst those of ESR1 with 151 out of 190 of them overlapping with those of ESR1 (hypergeometric *P* = 8e-239), consistent with molecular cross-talks between the two cancer biomarkers. The hormonal receptor genes ESR1 and PGR shared 22 MAGs (*P* = 1.9e-18), ESR1 and ERBB2 had 18 in common (*P* = 3.2e-27), ERBB2 and PGR had 3 (*P* = 0.0005), ERBB2 and TP53 had 21 (*P* = 4.5e-37) while TP53 and PGR had 16 genes in common (*P* = 5.3e-13; all P-values are estimates of enrichments of metastasis genes for one biomarker amongst that of the second biomarker taking into account the total number of genes in the network using a hypergeometric test). It is possible the overlap between metastasis genes across all these biomarkers could lead to regulatory cross-talks that enhance dysregulation of metastasis genes in tumors with simultaneous dysregulation of the TP53, ESR1, PGR and ERBB2, resulting in aggressive triple negative breast tumors.

**Figure 2.**
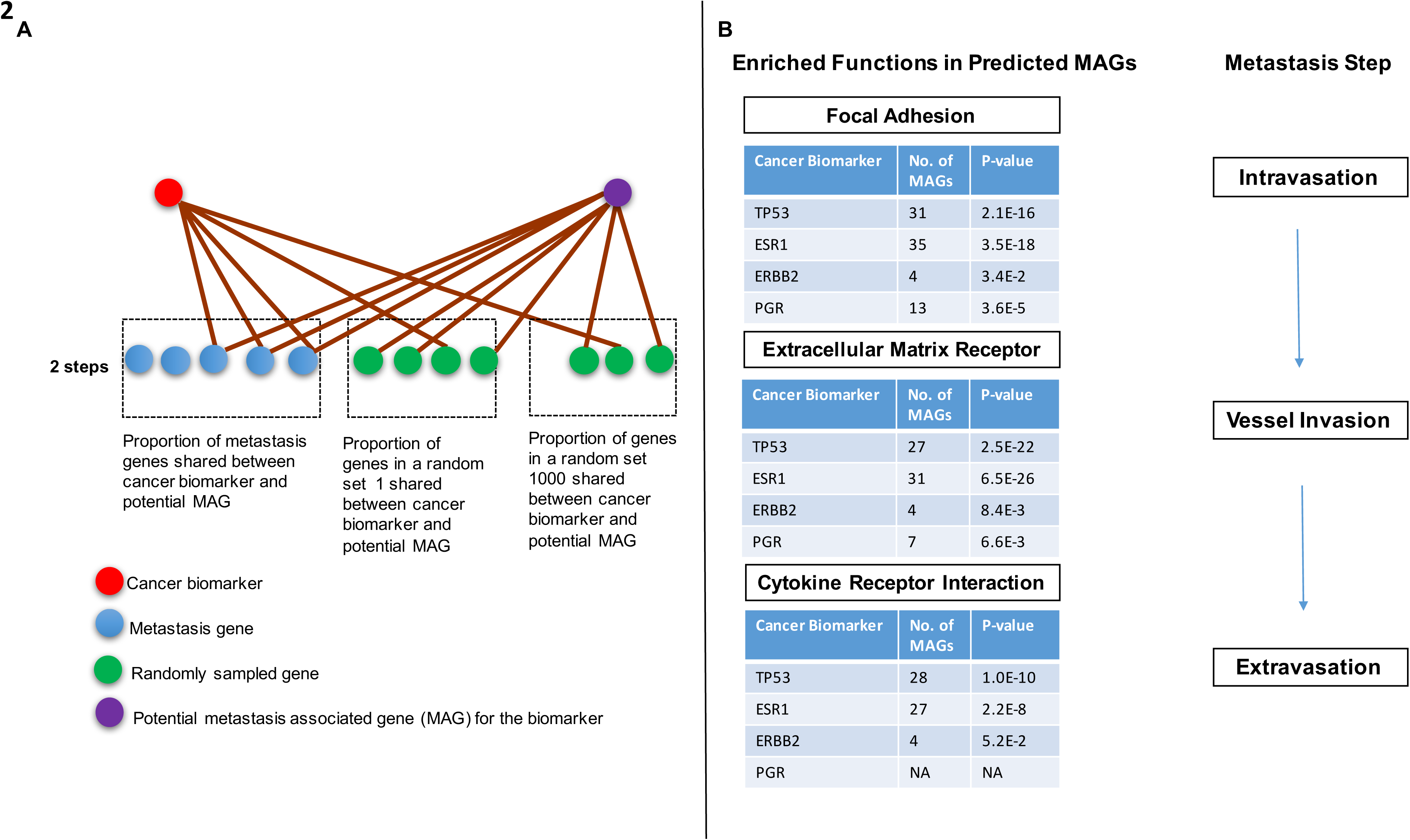
Prediction of Novel Metastasis Genes Associated with a Given Marker. (**A**) Schematic of the prediction approach. Depiction of topological relationships between TP53 and metastasis genes (red). Other genes (orange) that have a similar relationship to metastasis genes were identified in the network and are referred to as TP53 associated metastasis genes. Only genes whose proportion of 2^nd^ degree neighbors that are metastasis genes shared with TP53 are larger than any of the overlaps observed in 1000 sets of randomly sampled groups of genes were considered as TP53 associated metastasis genes (P < 0.001). (**B**) Summary of predicted metastasis genes associated with each biomarker and biological function enrichments.

#### Predicted MAGs are Associated with Metastasis in Breast Cancer Patients

Cancer involves both defects in cell division as well as the acquisition of new abilities that allow tumor cells to spread to distant tissues. Both processes affect prognosis of cancer patients- breast cancer prognostic signatures such as the 70-gene expression signature include genes involved in cell division as well as metastasis^51^. To validate the clinical relevance of the predictions, we next asked whether predicted metastasis genes are specifically associated with metastasis in patient derived data. Recently, a prognostic model for breast cancer based on highly co-expressed genes across multiple cancers (attractor metagenes) was demonstrated to outperform several models in predicting survival of about 2000 patients ^52,53^. We examined whether the top 10 genes in the attractor metagene associated with the transition of breast cancer from *in situ* carcinoma to invasiveness (mesenchymal or MES metagene) are predicted with high confidence by our approach as potential MAGs relative to a similar number of genes in the mitosis (CIN) and lymphocyte-specific immune recruitment (LYM) metagenes. For TP53-MAGs, 3 MES metagenes (SPARC, THBS2 and CTSK) were predicted with high confidence as metastasis associated (*P* < 0.001) and all but one MES gene had *P* > 0.05 (Supplementary Material). None of the genes in the mitotic metagene had *P* < 0.05 and only 2 genes in the LYM metagene had *P* < 0.05, demonstrating that the predicted TP53-associated metastasis genes are preferentially enriched with genes that correlate with metastasis relative to mitosis or immune recruitment. We observed similar results for the predicted ESR1, ERBB2 and PGR MAGs (Supplementary Material). The distinct enrichment of MAGs for functions involved in metastasis but not for those in cell division, in spite of the fact that the predictions are based on data from the MCF7 cell line that is poorly metastatic, validate the predictions.

The predicted MAGs that strongly correlate with metastasis in patient data (i.e present in MES metagene) may distinctly affect the occurrence of distant metastases in patients and potentially vary by TP53, ESR1, ERBB2 and PGR status. In other words, the impact of the metastasis genes in clinical metastases may differ between patients depending on their tumor biomarker status. Of the 10 genes in the MES metagene, 2 (SPARC and THBS2) were predicted at high confidence as MAGs (*P* < 0.001) with respect to more than one of the four biomarkers. Therefore, we examined further the correlation between these MAGs and distant metastases free survival (DMFS) in patients stratified by each of the biomarkers using public datasets^54^ (Methods). When considering all patients irrespective of their TP53 status, we found that expression levels of the SPARC gene was not associated with DMFS (*P* = 0.83; Figure 3 A). However, when patients were stratified by TP53 status, those with wild-type TP53 and high expression levels of SPARC had longer time to distant metastases compared to wild-type TP53 patients with low expression of the gene (*P* = 2.2e-05; Figure 3 B). In contrast, among patients with mutant TP53, those with low expression levels of SPARC were not different from those with high expression of the gene in the time to distant metastases (*P* = 0.78; Figure 3 C). Similarly, the expression level of SPARC in patients stratified by the other biomarkers (ESR1, ERBB2 or PGR) did not associate with DMFS (Supplementary Material). SPARC (secreted protein acidic and rich in cysteine) gene affects several metastasis associated processes including cell adhesion^55^ and remodeling of the extracellular matrix^56^. Its overexpression is associated with poor prognosis in multiple cancers including breast tumors^57^.

**Figure 3.**
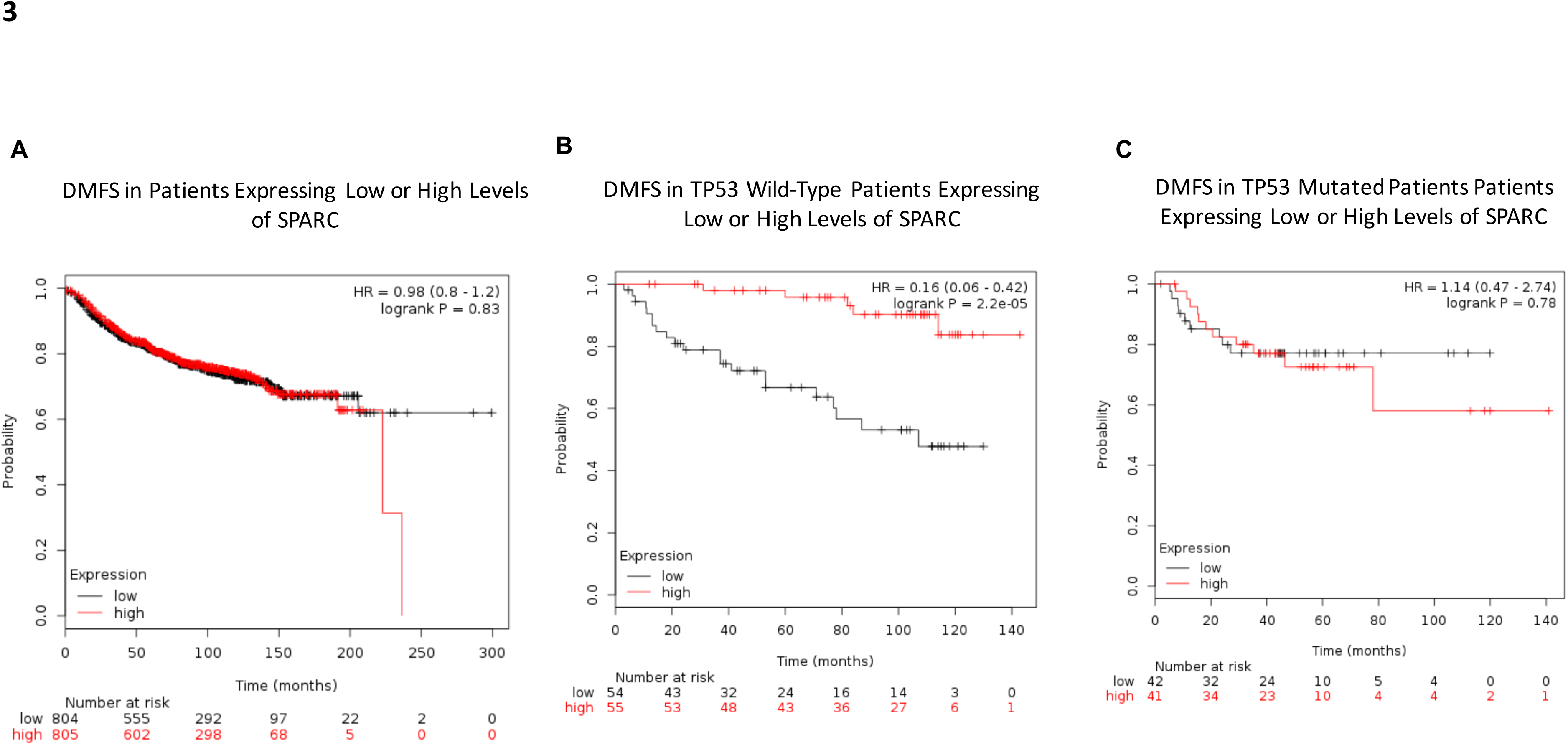
Context dependent associations between a metastasis gene and distant metastases.

Notably, SPARC and TP53 have inverse effects on apoptotic signaling and overexpression of SPARC reduces TP53 protein levels^58^, providing a plausible explanation of the observed TP53-dependent relationship between SPARC expression and DMFS.

Next, we examined the association between THBS2 expression and DMFS (Supplementary Material). In ESR1 positive patients, there was no difference in DMFS between patients based on expression of THBS2 (*P* = 0.75) but in the ESR1 negative group, patients expressing low levels of THBS2 had a longer time to DMFS compared to those expressing high levels (*P* = 0.026). There were no significant differences in DMFS based on THBS2 expression when patients were stratified by either TP53 or ERBB2 (Supplementary Material). Stratification of patients based on PGR revealed differences in DMFS between patients expressing low or high levels of THBS2 with the former group of patients showing a longer time to DMFS (*P* = 0.029), similar to the observation when stratification was based on ER status. The THBS2 promoter contains estrogen response elements which have been validated by chromatin immunoprecipitation and shown to mediate the ability of estrogen to suppress invasion by prostate cancer cells^59^. While both SPARC^56^ and THBS2^59^ have been implicated in breast cancer metastasis, our study predicted these genes using data from a cell line and raise the possibility that the impact of these genes on metastasis free survival could be dependent on breast cancer subtypes. These results imply that there could be value in personalized prognostic models based on breast cancer subtypes as the impact of specific genes on metastasis may vary across breast cancer subtypes. Similarly, therapeutics that target metastatic pathways could potentially have variable activity across patients depending on their tumor genetic background.

#### Identification of Master Regulators of Breast Cancer Metastasis

To identify potential regulators of MAGs for each biomarker, we queried chromatin immunoprecipitation datasets and miRNA targets using Enrichr^48^. MAGs for both TP53 and ESR1 were highly enriched with binding sites for the transcription factor BACH1, with this transcription factor showing the highest enrichment significance (*P* = 5e-15 for TP53-MAGs and *P* = 1.8e-24 for ESR1-MAGs; Supplementary Material). BACH1 is an experimentally validated master regulator of breast cancer metastasis to bones regulating the expression of several metastasis promoting genes including MMP1 and CXCR4, its ectopic expression enhances breast cancer malignancy and is associated with increased risk of breast cancer recurrence in patients^60^. Furthermore, a recent study reported that BACH1 promotes single cell heterogeneity between breast cancer cells leading to non-genetic variability and facilitating metastatic transitions in which a fraction of non-invasive cells are switched into a prometastatic state^61^. BACH1 has primarily been validated for metastases to bone, which are not only the most common metastases in breast cancer but also more frequent in estrogen receptor positive/ progesterone receptor negative tumors^26^. The high enrichment of many of the predicted ESR1-MAGs with BACH1 targets provides one possible mechanism for the role of ESR1 in bone metastasis. This observation opens the possibility of predicting organ specific metastases using the predicted MAGs that are specifically associated with one cancer biomarker and not others. The top regulators for PGR and ERBB2-MAGs were TCF4 and EGR1 (early growth response-1), respectively. BACH1 (see above), SOX2, EGR1 and RELA were among the top 5 master regulators for MAGs for more than one biomarker (Supplementary Material). SOX2 was in the top 5 master regulators of MAGs for all the four biomarkers (for TP53 metastasis genes *P* = 2.8e-12; for ESR1 *P* = 3.8e-14; for ERBB2 *P* = 0.0008; for PGR *P* = 8e-11; and Supplementary Material), in agreement with its key role in tumor initiation and cancer stem cell functions^22^ with the latter playing a pivotal role in metastasis^62^. EGR1 targets were significantly enriched with MAGs for ESR1 (*P* = 3.8e-14), PGR (*P* = 3.9e-13) and ERBB2 (*P* = 0.0002) while RELA targets were enriched with TP53-MAGs (*P* = 7.8e-15) and ESR1 (*P* = 9.4e-18), consistent with previous reports on the role of RELA^46^ and EGR1^63^ in metastasis. These results suggest that metastasis could be regulated through 2 main groups of transcription factors- those that are master regulators of molecular functions that are broadly important in metastases to different organs (for example, cancer stem cell functions) and those that facilitate metastases to specific organs. We provide a ranked list of the top 5 predicted master regulators in the Supplementary Material.

Next, we examined enrichments of the predicted metastasis genes for miRNA targets based on TargetScan and Enrichr ^48^. The top enrichments for miRNA targets were mir-342 for TP53-MAGs (*P* = 0.008), mir-29 for ESR1 (*P* = 0.007) and mir-30 for PGR (*P* = 0.0008; see Supplementary Material for ranked lists of miRNAs). In an analysis of the expression of 453 miRNAs in breast cancer subtypes, mir-342 was reported as highly expressed in ESR1 and ERBB2 positive luminal B tumors and downregulated in basal ones^64^. However, Lowery et al did not find associations between mir-342 and tumor stage or nodal status, although in colorectal cancer this miRNA inhibits proliferation and invasion^65^. mir-29 and mir-150 targets were enriched in the TP53 (*P* = 0.03) and ESR1-MAGs (*P* = 0.007; Supplementary Material). Both mir-29b and mir-150 directly target TP53 and have been linked to tumor malignancy^50,66^. On the other hand, mir-29b regulates epithelial to mesenchymal transition, an important step in metastasis^67^. mir-150 has been validated as promoting breast tumor growth and malignancy^68^ while mir-30 regulates non-attachment growth of breast cancer cells^69^. ERBB2 metastasis associated genes were not enriched for any of the miRNA targets at a P-value threshold of 0.05.

#### Predicted Metastasis Genes and Organ Specific Tropism in Breast Cancer Subtypes

To understand the potential role of the predicted metastasis genes in organ specific metastasis, we determined overlaps between the predicted MAGs for each biomarker and published differentially expressed genes associated with metastases to bones^32^, lungs^34^ and brain^33^. The top enrichment for TP53 associated metastasis genes was for brain specific metastasis genes (HBEGF and COL13A1; *P* = 0.001; Supplementary Material) followed by an enrichment for lung specific metastasis genes (EREG and LTBP1; *P* = 0.04; Supplementary Material). Bone specific metastasis genes were not enriched among the TP53 metastasis genes (*P* = 0.28). In contrast, they were the top enriched for ESR1-MAGs (CTGF, FYN and FST; *P* = 0.01), followed by those of the brain (HBEGF and COL13A1; *P* = 0.03) and lungs (EREG and VCAM1; *P* = 0.04). There were no significant enrichments for the ERBB2 and PGR predicted MAGs for organ specific metastasis genes. The enrichment of TP53-MAGS for genes associated with brain metastases is consistent with an increased risk of brain metastases in patients with TP53 mutations^11,42^ while the enrichment of bone associated metastasis genes in the ESR1-MAGs is consistent with high incidence of bone metastases in estrogen receptor positive patients^12,26^. Though ERBB2 has also been associated with increased risk of brain metastases^10^, its predicted MAGs did not include any of the genes associated with brain metastasis, potentially because the MCF7 cell line expresses normal level of ERBB2 while breast tumors with increased risk to brain metastases have high ERBB2 expression levels.

To further validate the association between the predicted MAGS and organ specific metastases, we compared them to ECM components that have been associated with organ specific metastases^70,71^. We determined overlaps between the predicted MAGs and ECM integrins reported to constitute *in vitro* phenotypic fingerprints that are predictive of *in vivo* organ specific metastases^71^ (Figure 4 A) Integrin alpha 2 (ITGA2) which is associated with brain/ lung metastases^71^ was present only among TP53-MAGs (*P* = 0.05) but not in ESR1-MAGs. Both TP53 and ESR1-MAGs included at least 1 integrin associated with binding to lung or bone ECM (Figure 4 A). Specifically, TP53-MAGs included ITGA3 (associated with bone metastasis, *P* = 0.21) as well as ITGA1, ITGA2 and ITGA10 (the latter 3 associated with lung metastasis, *P* = 0.001). ESR1-MAGs included 3 integrins associated with metastases to the lungs (ITGA1, ITGA5 and ITGA10; *P* = 0.0007) alongside 3 other integrins associated with bone metastases (ITGA3, ITGA5 and ITGB7; *P* = 0.0007) but did not include the brain metastasis integrin, ITGA2 (*P* = 0.95; Figure 4 A; all P-values based on hypergeometric test taking into account the total number of metastasis genes for a given biomarker, the number of integrins associated with the given tissue and the number of integrins associated with the tissue that are also present in the list of the metastasis genes for the given biomarker). ERBB2-MAGs did not include any of the bone (*P* = 0.6) or brain integrins associated with metastasis (*P* = 0.9) and had only 1 lung associated integrin (ITGA10, *P* = 0.4). Thus, compared to the other biomarkers, TP53-MAGs had the highest enrichment for brain metastasis genes while ESR1 had the highest enrichments for bone and lungs consistent with breast cancer metastasis patterns in patients with aberrations in these biomarkers^11,12,26,42,59^.

**Figure 4.**
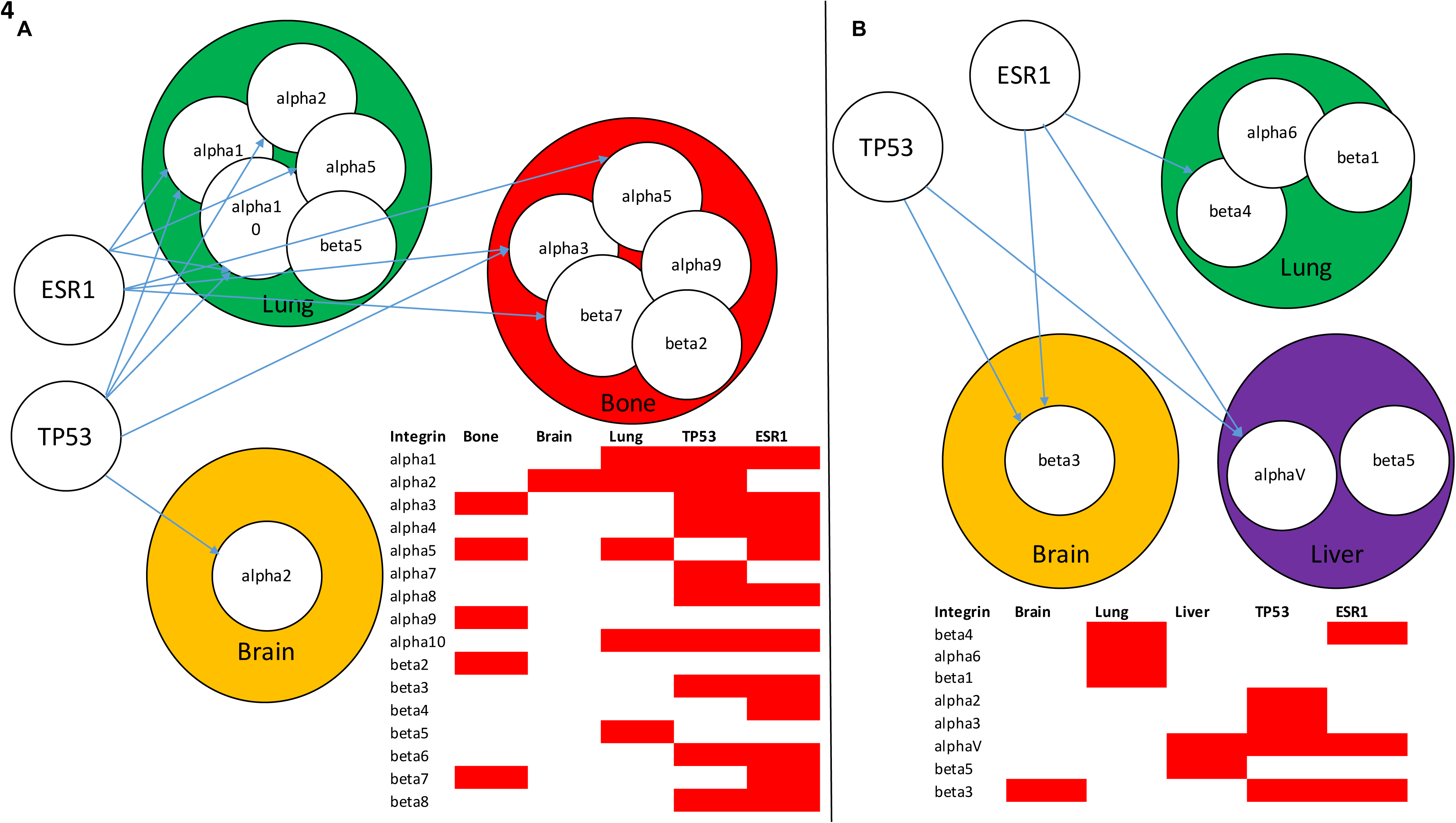
Different cancer biomarkers may promote metastasis to specific organs due to molecular relationships with pathways associated with specific organs, here the different ECM components. Connections between metastasis genes for TP53 and ESR1 with (**A**) integrins mediating organ specific metastases identified using a synthetic biomaterial predictive of organ specific and, (**B**) connections between metastasis genes for TP53 and ESR1 with exosome derived integrins mediating organ specific metastases.

Recently, exosome derived integrins were found to predict metastasis to the brain, bone and lungs^70^. Therefore, we repeated the analysis above using the exosome derived integrins to determine their enrichment in MAGs for each biomarker (Figure 4 B). TP53-MAGs included ITGB3 (*P* = 0.02), an integrin whose presence in tumor derived exosomes was associated with increased metastasis to the brain and ITGAV (*P* = 0.04, Figure 4B) which was associated with metastasis to the liver^70^. ESR1-MAGs included integrins associated with metastasis to the brain (ITGB3, *P* = 0.01), liver (ITGAV, *P* = 0.02) and lungs (ITGB4, *P* = 0.02). ERBB2-MAGs did not include any of the exosome derived integrins while PGR-MAGs had only 1 exosome derived integrin (ITGAV, *P* = 0.02), associated with liver metastases (Supplementary Material). Besides integrins, other organ specific components such as collagens in the ECM could influence metastasis. We provide a full comparison of the ECM components specific to TP53-MAGs vs. those of ESR1-MAGs in the Supplementary Material.

In summary, it appears that each biomarker is a short distance to molecular networks of genes associated with metastasis to more than one organ even though the strength of these connections could be stronger for some organs vs. others and vary across biomarkers and metastasis pathways. Potentially, combining multiple levels of evidence for the association between a given biomarker and metastasis to specific organs may provide a more accurate overall likelihood for metastases to specific organs. Consequently, TP53-MAGs exhibited the highest association with brain associated metastasis genes derived from 3 sources: i) brain specific metastasis genes^33^, ii) integrins identified using *in vitro* phenotypes^71^ and iii) exosome derived integrins^70^. ESR1-MAGs had the highest association to bone metastasis genes, consistent with increased frequency of bone metastasis in estrogen receptor positive patients^12,26,72^ in contrast to increased risk for visceral metastases in patients with aberrations in TP53 ^11,42,72,73^. Brain metastases have worse prognosis, neurological complications, low quality life, short survival and are more difficult to treat as many drugs are unable to cross the blood brain barrier^14,74^. The strong association between TP53 and brain associated metastasis genes could enable future understanding of the role of TP53 in this process especially given that mutations in the gene are also associated with aggressive breast tumors^19^. Collectively, these results suggest that different breast cancer subtypes may utilize distinct molecular routes to metastasis, even to the same organ. Thus, knowledge of these subtype specific routes to metastasis could aid in discovery of personalizing therapeutics.

#### Predicted Metastasis Genes are enriched with Perturbation Signatures of Agents Influencing Metastasis

Chemical agents that alter the expression of the predicted MAGs could potentially influence metastasis and thus predict potential therapeutic targets for metastasis. While such a result may be deduced from the observation that MAGs are enriched with genes that influence metastasis, unexpected context specific effects of metastasis blocking agents could arise as distinct groups of co-regulated MAGs may exist. That is, the expression of some groups of MAGs may be affected by one metastasis inhibiting agent but not another. To test this, we queried gene expression signatures of cancer cell lines exposed to thousands of drugs (Enrichr/ LINCS1000 database^48^, see Methods) to determine perturbations whose signatures are enriched with MAGs. Gene expression changes induced by perturbations by several inhibitors of cancer-related kinases including focal adhesion kinase (FAK), EGFR, MEK and BRAF were the most enriched with predicted MAGs for TP53, ESR1, PGR and ERBB2 (Supplementary Material). Interestingly, genes upregulated by FAK inhibitors were the most enriched in TP53-MAGs (*P* = 8.7e-11), ESR1-MAGs (*P* = 6.9e-10) and ERBB2-MAGs (*P* = 7.5e-8, Supplementary Material), consistent with the role of FAK in several processes that enhance metastasis including cell migration, invasion, survival and cancer-stem cell self-renewal^20^. PGR-MAGs were most enriched among genes whose expression was upregulated by the phosphoinositide-3-kinase (PI3K) inhibitor, PI-103 (*P* = 2e-5, Supplementary Material). In contrast, genes downregulated by the investigational PI3K inhibitor GSK-1059615^75^ (*P* = 4.4e-8) were the top enriched with TP53-MAGs. This result is supported by the roles of PI3K in cancer: the gene encoding its catalytic subunit PI3KCA is the second most frequently mutated gene across multiple cancers^30^, and is involved in nearly all human tumors through its effects on growth factor signaling, cell proliferation, metabolism and survival^76^. PI3K has also been associated with metastasis^77^ and its inhibition can paradoxically enhance metastasis^78^. This observation suggests the possible metastatic effect that drugs primarily considered as interfering with growth related processes could have. It is further supported by the observation that differentially expressed genes in perturbations by drugs inhibiting EGFR (afatinib, clinically approved and pelinitib) showed an enrichment in MAGs (Supplementary Material). Surprisingly, only TP53-MAGs were enriched with differentially expressed genes of angiogenesis inhibitors that target the vascular endothelial growth factor receptor (VEGFR) or platelet derived growth factor receptor (PDGFR). Angiogenesis-the formation of new blood vessels from existing ones-enhances growth and metastasis of cancer cells by providing tumor cells with increased oxygen and nutrient supply^79,80^. Genes downregulated by perturbations of the VEGFR/ PDGFR inhibitor linifanib were enriched with TP53-MAGs (*P* = 5e-6, Supplementary Material). Interestingly, TP53 inhibits angiogenesis^81^ and it has been reported that the effect of angiogenic therapy in patients is sensitive to TP53 status^82^. Thus, biomarker specific MAGs may be associated with variations in patient outcomes to metastasis blocking therapeutics. Notably, some of the drugs whose perturbations were enriched with the metastasis genes are under investigation for other non-cancer indications. For example, genes downregulatd by perturbations of oxibendazole-an anti-helminthic compound^83^ or QL-XII-47- an investigational compound for dengue virus infection^84^ were enriched with PGR-MAGs (*P* ∼ 0.0004; Supplementary Material). Hence, drugs for other diseases could potentially be repositioned for metastasis. We provide a table in the Supplementary Material of the top 10 perturbations that were enriched with up- or downregulated genes for MAGs for each biomarker.

#### Population Differentiation of Predicted Metastasis Genes

It is known that breast cancer progression differs across populations due to a wide range of factors including socio-economic and biological factors^85–88^. Population genetic variation in candidate genes associated with metastasis could potentially contribute to population disparities in cancer progression, and lead to variations in the effectiveness of metastasis targeting therapeutics. Therefore, we obtained fifteen positive selection measures for the predicted MAGs (Supplementary Table 4) using recent positive selection measures across human populations in the 1000 genomes ^89^ and HapMap II^90^ projects (dbPSHP database)^91^. For ERBB2-MAGs, the derived allele frequencies (DAF) of SNPs in EPAS1, COL14A1 and LAMC2 showed the highest rates of positive selection when comparing European (CEU, Central Eastern European) and African (YRI, Yoruba) populations (for detailed statistics see Supplementary Table 4). Recently, EPAS1-a gene involved in response to hypoxia and adaptation to high altitudes-emerged as one under the highest signature of positive selection in Tibetans^92–94^. In relation to this, tumor hypoxia is a risk factor for metastasis, predicts poor outcomes in patients^95,96^, and is associated with activation of genes that respond to low oxygen concentrations, for example, EPAS1 (hypoxia inducible factor-2 alpha) ^97^. ERBB2 signaling increases the expression of hypoxia inducible factor 1 alpha (HIF1-alpha)^98^ and EPAS1 expression correlates with hypoxia-related gene expression in ERBB2+ breast cancers^99^. It is therefore notable that EPAS1 was predicted specifically with high confidence as metastasis associated for ERBB2 (*P* < 0.001) as compared to its relationship with the other biomarkers (for TP53 *P* = 0.009, ESR1 *P* = 0.003, PGR *P* = 0.005). Moreover, we found that EPAS1 expression is significantly associated with DMFS in ERBB2+ (*P* = 0.0087, Figure 5) but not ERBB2- breast cancer patients (*P* = 0.48, Figure 5). The other 2 metastasis genes with high DAF (COL14A1 and LAMC2) have been associated with metastasis in renal cancer^100^ and lung adenocarcinoma^101^, though we found no associations to metastasis in breast cancer using literature. In addition to the 3 genes above with high DAF, another ERBB2-MAG with high population specific selection is the Duffy antigen receptor for chemokines (DARC), a gene that is widely known for its association with resistance to malaria in African populations^102,103^. Interestingly, increased expression of DARC inhibits growth and metastasis of breast cancer cells potentially by sequestering angiogenic cytokines^104^. The Duffy Blood Group Antigen also correlates with breast cancer incidence, lymph node metastasis and overall survival^105^. According to our analysis, in ERBB2+ breast cancer patients, there is no association between DARC expression and metastasis (*P* = 0.17, Figure 5) while in ERBB2- patients, high DARC expression is significantly associated with delayed time to distant metastases (Figure 5, *P* = 0.0092). Therefore, while DARC negative individuals (highly frequent in individuals of Africa descent), may have protection from malaria, they may be more prone to distant metastases, especially in African women (ERBB2-). Furthermore, triple negative status (ESR1-/PGR-/ERBB2-) is also higher amongst African women^106^ which is associated with low responsiveness to hormonal and HER2-targeted therapies such as tamoxifen and Herceptin, respectively. Thus, understanding why DARC expression is associated with better prognosis especially in ERBB2-women may help identify therapeutics for this group. We provide population differentiation measures for each of the predicted metastasis genes in Supplementary Table 4. These results also point to the conflicting effects of genetic variants that confer protection from one environmental condition or disease while increasing susceptibility or severity to other unrelated conditions, diseases or pharmacological agents.

**Figure 5.**
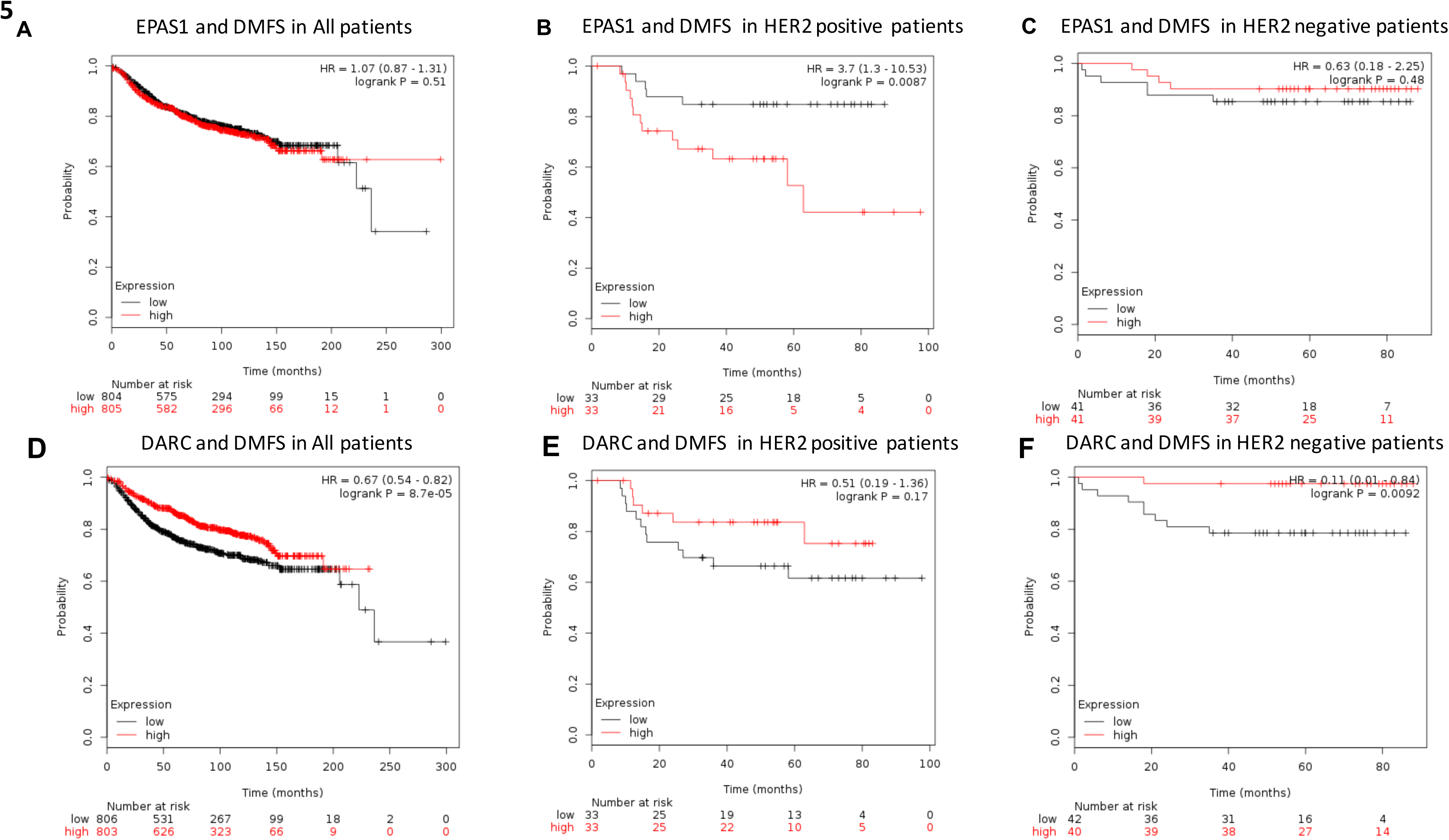
Association between distant metastases and expression of 2 metastasis associated genes showing high population differentiation in African vs. European populations.

## Discussion

This study has generated a number of novel insights on breast cancer metastasis: i) the major cancer biomarkers (ESR1, PGR and ERBB2) used to classify breast cancer into molecular subtypes alongside TP53-a gene involved in multiple cancers-are a short path from the molecular networks of metastasis genes. ii) By leveraging this unique topological association, we have shown that genes sharing this property (MAGs) with the biomarkers are enriched with functions that are key to metastasis. We have validated the clinical significance of these genes by demonstrating their association with distant metastases in breast cancer patients. iii) The predicted MAGs are potential targets for metastasis blocking therapeutics and may be important in personalizing therapy. vi) We found a strong association between TP53-MAGs and signatures of brain metastases compared to the strong association between ESR1-MAGs and bone metastases, suggesting that the approach developed in this study could be used to predict organ specific metastasis. iv) Some MAGs have been under previous positive selection in different human populations and may be therefore associated with population differences in breast cancer prognosis and therapeutics. In particular, the observation that low expression of the malaria resistance gene DARC^103^ is associated with increased risk of distant metastases especially in ERBB2 negative patients (or ESR1 positive patients, Supplementary Material) has high clinical relevance as this group of patients have lower benefit to herceptin. In addition, African women are more likely to express overall low levels of DARC (due to a DARC negative phenotype on their red blood cells) and epidemiological correlations between breast cancer and DARC alleles have been reported^105^. Therefore, these observation may be leveraged for the development of targeted therapeutics for this group of patients who are also more prone to aggressive breast tumors^87^. Thus, the findings in this study could enable basic understanding of metastasis as well as aid in the development of clinical interventions for predicting and blocking the process.

A potential drawback of our study is that it is based on an interactome obtained from the MCF7 cancer cell line. The MCF7 cell line was derived from an invasive breast ductal carcinoma of a Caucasian woman^107^. It is ESR1 and PGR positive, HER2 (ERBB2) negative and contains wild-type TP53^108,109^. MCF7 is poorly metastatic but can be induced to a metastatic state by estradiol and is one of the most widely used cancer cell lines^110^. In spite of this drawback, the predicted metastasis genes are consistent with well-known processes in metastasis as shown by a high enrichment of ECM components, focal adhesion, cytokine-cytokine interactions and chemokine signaling. In addition, we show that the predicted genes are specifically enriched with genes associated with metastasis in breast cancer patients and can discriminate them from genes involved in cell division which play key roles in tumor initiation. Our results also shows that the transcript levels of some of these genes are associated with distant metastasis free survival (DMFS) and that these associations can differ between groups of patients stratified by their tumor biomarker status. It is possible that the reason only a few ERBB2-MAGs were predicted and no association between ERBB2 and organ specific metastasis was found is because of the low ERBB2 expression in the MCF7 cell line compared to ERBB2 positive tumors. Another limitation of the study is that we still have limited knowledge on the specific genes required for organ specific metastasis and survival of tumor cells in distinct tissues. Increasing functional validation of organ specific mediators of metastasis will help increase the accuracy of the predictions for organ specific metastasis at an individualized patient level. In spite of this, we demonstrate that the limited set of organ specific metastasis genes when combined with our approach predicts strong associations between ESR1 and bone metastases, and TP53 and visceral metastases, consistent with epidemiological observations^11,12,26,28^. We have addressed in part some of these limitations by validating our predictions on patient derived data sets, several other independent studies that identified metastasis genes or biological pathways required for metastasis and studies on organ tropic behavior of breast cancer subtypes.

Our study highlights the possibility that tumor biomarkers may potentially influence metastasis trajectories in distinct ways due their closeness to distinct biological processes associated with metastasis, potentially leading to increased risk of metastases to certain organs and causing inter-patient variations in response to metastasis blocking agents. In future, the approach outlined here can be applied to molecular networks derived from other cancer cell lines or even patient derived gene interactomes, potentially paving way for targeted or personalized interventions to metastases of various tumors.

## Materials and Methods

### Breast Cancer Interactome

The breast cancer interactome was obtained from the Califano Lab at Columbia Univerisity Center for Multiscale Analysis of Genomic and Cellular Networks, MAGNet (http://wiki.c2b2.columbia.edu/califanolab/index.php/Califano_Info). The MCF7 interactome was constructed by applying the reverse engineering algorithm ARACNe to 3000 gene expression profiles of MCF7 breast cancer cell lines exposed to 448 drugs in replicates and at various doses (CMAP2 dataset).

### Selection of Metastasis Associated Genes from Literature

We compiled a list of genes from a review of protein based biomarkers associated with metastasis of various cancers including ovarian, breast, colorectal, lung, endometrial, pancreatic, cervical, prostrate, gastric and others based on several previous studies^36^. We selected validated protein based biomarkers because of their high translatability to the clinic. The biomarkers were previously discovered and validated using various approaches including mass spectrometry (iTRAQ, LC-MS), ELISA, phage display, 2DIGE, RT-PCR, immunohistochemistry.

### Assessment of Molecular Associations between Cancer Biomarkers and Metastasis Genes

For each biomarker, the number of known metastasis genes located 1, 2, 3,4 and 5 steps away from its node in the MCF7 interactome was determined as follows using the R package igraph. First, pairwise distances (path length) between all genes in the network were determined using the function distances in igraph. The resulting matrix of pairwise distances between genes was then queried for distances between each biomarker and the metastasis gene set from literature. The number of metastasis genes located at n steps (1, 2, 3, 4 and 5 path length) were then determined for each biomarker. Next, 1000 randomly sampled gene sets, where each gene set contained the same number of genes as the metastasis gene set (35 genes), were sampled. The matrix of pairwise distances was then queried to determine the number of genes in each gene set that are n steps away from the biomarker of interest. For each biomarker, the number of metastasis genes, m, located n steps away from its node in the network was compared to the number of genes in each random gene set that are n steps away from the biomarker. The likelihood of observing m genes that are n steps away from the biomarker by chance was estimated as the proportion of the 1000 random gene sets in which the count of genes that are n steps away from the biomarker is greater than that of m genes that are n steps away from the biomarker.

### Prediction of Metastasis Associated Genes (MAGS)

For each gene, *g_i_*, in the genome, we first identified metastasis genes that are its 2^nd^ degree neighbors. Then, we computed the fraction of these genes that overlap with 2^nd^ degree neighbors of the biomarker and compared this to that observed in 1000 sets of randomly sampled genes (Figure 3 A and Methods). To determine the statistical significance of the overlap, we randomly selected 35 genes from the genome and computed the proportion of these genes that are 2^nd^ degree neighbors of both the biomarker and the gene, *g_i_*. We repeated this process 1000 times to obtain a distribution of the fraction of genes in 1000 random sets of 35 genes that happen to be common 2^nd^ degree neighbors of a given biomarker and a given gene. We considered *g_i_* to be a potential metastasis associated gene if the overlap between metastasis genes that are its 2^nd^ degree neighbors as well as a 2^nd^ degree neighbor of the biomarker was greater than the overlap observed in each of the 1000 random instances (P < 0.001). We also obtained q-values or FDR adjusted P-values using the SVA package in R (p.adjust function and the Benjamini Hochberg method).

### Construction of Survival Curves for Distant Metastasis Free Survival

Survival curves were obtained using an online tool – the Kaplan Meir Plotter (http://kmplot.com/analysis/index.php?p=service&cancer=breast) for assessing the effect of expression of 54, 675 genes on survival using a compilation of 5, 143 breast cancer samples. The survival analysis was done using the 2014 version of the Kmplot online system comprising 4142 breast cancer samples. Only jet-set probes were considered in the analysis for selecting the optimal microarray probeset for a representing genes.

